# The probability of joint monophyly of all species in an arbitrary species tree

**DOI:** 10.1101/2021.12.16.473006

**Authors:** Rohan S. Mehta, Mike Steel, Noah A. Rosenberg

## Abstract

Monophyly is a feature of a set of genetic lineages in which every lineage in the set is more closely related to all other members of the set than it is to any lineage outside the set. Multiple sets of lineages that are separately monophyletic are said to be reciprocally monophyletic, or jointly monophyletic. The prevalence of reciprocal monophyly, or joint monophyly, has been used to evaluate phylogenetic and phylogeographic hypotheses, as well as to delimit species. These applications often make use of a probability of joint monophyly under models of gene lineage evolution. Studies in coalescent theory have computed this joint monophyly probability for small numbers of separate groups in arbitrary species trees, and for arbitrary numbers of separate groups in trivial species trees. Here, generalizing existing results on monophyly probabilities under the multispecies coalescent, we derive the probability of joint monophyly for *arbitrary* numbers of separate groups in *arbitrary* species trees. We illustrate how our result collapses to previously examined cases. We also study the effect of tree height, sample size, and number of species on the probability of joint monophyly. The result also enables computation of relatively simple lower and upper bounds on the joint monophyly probability. Our results expand the scope of joint monophyly calculations beyond small numbers of species, subsuming past formulas that have been used in simpler cases.

## 1 Introduction

Evaluations of the prevalence of reciprocal, or joint, monophyly in sampled gene genealogies have been useful in a variety of studies in phylogenetics, phylogeography, and molecular ecology. They have been used for identifying units for conservation [Moritz, 1994], analyzing differing phylogeographic patterns across species [Carstens and Richards, 2007], evaluating the distinctiveness of taxa [Kubatko et al., 2011], and providing context for estimation of species divergence times [Arbogast et al., 2002]. Joint monophyly is fundamental to genealogical perspectives on species delimitation [Hudson and Coyne, 2002, De Queiroz, 2007].

Central to the application of joint monophyly is a theoretical prediction of the probability that genealogies show joint monophyly as a function of evolutionary parameters. Many studies have used monophyly probability computations in studies of the evolutionary relationships among recently-diverged species [Birky et al., 2005, Carstens and Knowles, 2007, Carstens and Richards, 2007, Syring et al., 2007, Jansen et al., 2010, Kubatko et al., 2011, Bergsten et al., 2012, Rabeling et al., 2014]. These computations primarily made use of theoretical results of Rosenberg [2003, 2007], which consider the probability that gene lineages in two populations of species are jointly monophyletic as a function of population divergence times. For example, Kubatko et al. [2011] used such computations to assess the taxonomic distinctiveness of two species of *Sistrurus* rattlesnake, each of which was divided into three subspecies. They considered two types of comparisons for each set of three subspecies: first, that one subspecies was distinct from a hypothetical clade containing the other two, and next, that the two remaining subspecies were distinct from each other. The result of these comparisons was the establishment of the distinctiveness of a seriously threatened subspecies (*S. catenatus catenatus*), as well as varying levels of distinctiveness among the remaining subspecies.

Because the probability formulas available were limited to two groups, Kubatko et al. [2011] were restricted to performing a hierarchical set of analyses in which distinctiveness of one subspecies from a taxon that combined the other two subspecies was assessed, followed by distinctiveness of one of the two previously-combined taxa from the other. Joint monophyly computations were likewise restricted to these two hierarchical pairs of subspecies. Although this hierarchical analysis did produce the desired determinations, the analysis of Kubatko et al. [2011] would have been enriched by the ability to simultaneously consider the distinctiveness of one *S. catenatus* subspecies from the two other *S. catenatus* subspecies, rather than being restricted to a hierarchical pairwise comparison that might produce inaccurate probabilities as a result of merging present-day samples from populations that have diverged in the past [Mehta et al., 2016]. Furthermore, simultaneously studying the relationship between the *S. catenatus* subspecies in relation to the other *Sistrurus* species would have required results that were able to accommodate up to six simultaneous monophyly events. Other similar studies involving more than two species or groups have also been restricted to pairwise computations [Carstens and Richards, 2007, Baker et al., 2009, Neilson and Stepien, 2009, Kubatko et al., 2011, Bergsten et al., 2012].

Three theoretical developments now place the possibility of a joint monophyly probability computation within reach for taxa related according to an arbitrary species tree. First, Zhu et al. [2011] computed the probability of joint monophyly for an arbitrary number of groups for lineages originating within a single population rather than evolving on a species tree. Next, Mehta et al. [2016] found the probability of joint monophyly in a species tree of arbitrary size, considering two classes of lineages. Finally, Mehta and Rosenberg [2019] found the full probability of joint monophyly for the lineages of species evolving on species trees with three or four species. Thus, the first extension generalized to an arbitrary number of groups whose lineages must be jointly monophyletic. The second produced an algorithm that allows for an arbitrary species tree. The third provided the simplest cases for a synthesis of the other two extensions.

In this study, we obtain the complete generalization: the probability of joint monophyly for an arbitrary number of groups in an arbitrary species tree. Figure 1 illustrates the results of Zhu et al. [2011] and Mehta et al. [2016] and how they relate to our computation. We study the effect of species tree parameters, such as tree height and sample size, on this probability. Because the result is computationally intensive, we provide lower and upper bounds on the probability of joint monophyly, as well as an alternative, potentially faster, method for numerical computation. Finally, we provide software that encodes the new formulas.

**Figure 1:**
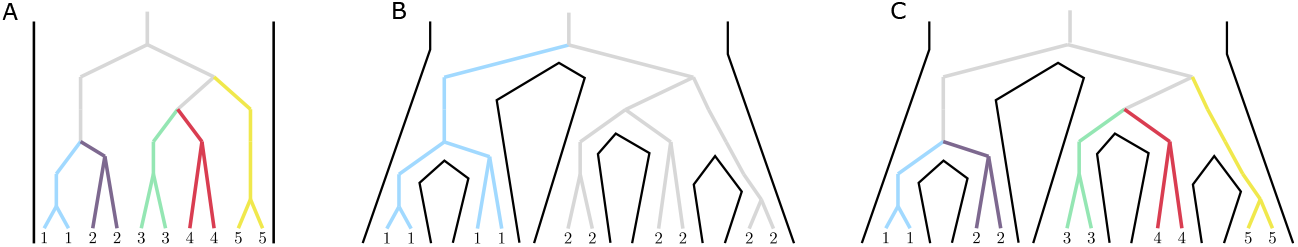
Schematic of the general joint monophyly calculation. (A) Zhu et al. [2011] computed the probability of joint monophyly of arbitrarily many groups in a single population. (B) Mehta et al. [2016] computed the probability of joint monophyly of two groups in an arbitrary species tree. (C) Here, we compute the probability of joint monophyly of arbitrarily many groups in an arbitrary species tree. In each panel, the numbers and colors indicate groups, and the black lines represent a species tree.

## 2 Preliminaries for the recursive approach

### 2.1 Model and notation

We consider a binary species tree *ℱ* on the species label set *S*, consisting of a topology and a set of branch lengths. For each leaf *S*_*i*_ of *ℱ*, we specify a sample size *s*_*i*_ ≥ 1. We use the multispecies coalescent to track the sampled lineages as they travel back in time “up” the species tree. Section 2 describes the terminology and construction of our coalescent model, closely following Mehta et al. [2016] and Mehta and Rosenberg [2019]. Figure 2 illustrates some of the notation.

**Figure 2:**
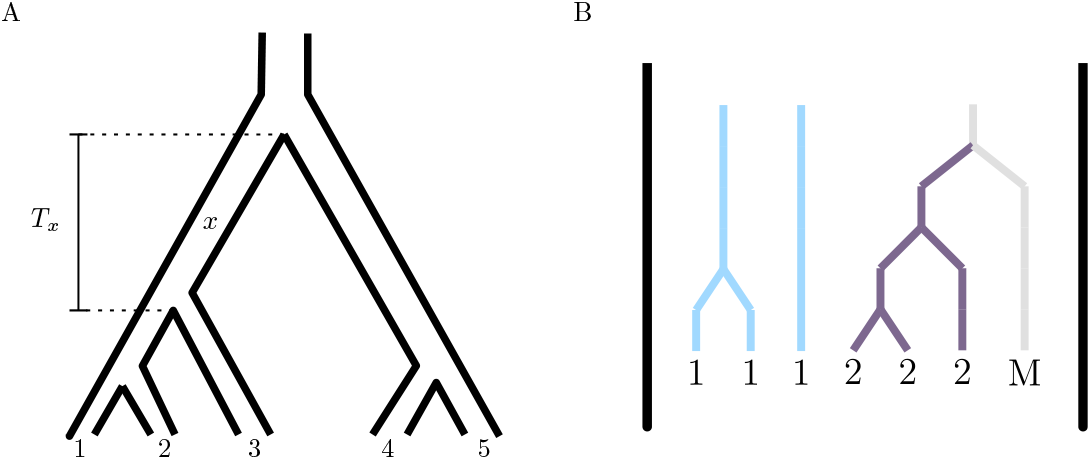
Notation for input and output lineages. (A) An example of a species tree *ℱ*, with five species and species label set *S* = *{*1, 2, 3, 4, 5*}*. An example branch *x* is highlighted with its branch length *T*_*x*_. (B) Coalescences happening within a single branch (branch *x* in (A)) of a species tree. In this diagram, three lineages from species 1, three lineages from species 2, and a single mixed lineage enter the branch, and two lineages from species 1 and one mixed lineage exit the branch. Supposing this branch comes from a five-species tree, the input state is 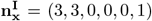, and the output state is 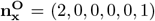. The label 1 is a surviving label, and the label 2 is a lost label.

### 2.2 Lineage labels

Genetic lineages are labeled according to the species from which they are sampled. All lineages for a particular species have the same label, and each species has a unique label. We label the species 1, 2, …, *k*, where the number of species is |*S*| = *k*. Lineages that result from a coalescence between lineages of differing labels are called “mixed” lineages and are assigned label *k* + 1.

### 2.3 Species tree branches

In our coalescent framework, the bottom of the tree is the present, at time 0, and time increases up the tree, further into the past. Viewed backward in time, an internal node of the species tree represents an event at which two species merge into an ancestral species. Gene lineages enter species tree nodes from the bottom and exit them the top as time progresses into the past. Because a one-to-one correspondence exists between species tree branches and nodes, we refer to a node and its immediate ancestral branch interchangeably. A particular node *x* has lineages enter from both branches directly below it. The length of branch *x* is *T*_*x*_, the time associated with node *x. T*_*x*_ is measured in units of *N* generations, where *N* is the haploid population size on branch *x*; this size is assumed to be constant over all species tree branches. Larger sizes correspond to smaller values of *T*_*x*_ in coalescent units. The root branch of *ℱ* is assumed to contain any coalescence events that have not occurred below the root. Biologically, this assumption is that of a universal common ancestor for all gene lineages, and it is implemented by setting the root branch length to infinity.

### 2.4 Input and output states

An output state of a branch *x* is a list of nonnegative integers that records the numbers of lineages of each label exiting the branch from the top. In our model, the output state is a random variable. This random variable is a vector *Z*_*x*_ of length *k* + 1 whose *i*th element is the number of output lineages that possess label *i*. A particular instance of this random variable is denoted 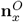.

Similarly, an input state for a branch is a list of nonnegative integers that records the numbers of lineages of each label entering the node from the two branches immediately below it. The input state for a branch *x* is the sum of the two output states for its descendant branches *x*_*L*_ and *x*_*R*_. A particular instance of an input state is 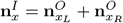. Figure 2B displays an example species tree node with its inputs and outputs.

### 2.5 Coalescence sequences

A coalescence sequence is an ordered sequence of coalescence events. For example, consider five lineages A, B, C, D, and E. One possible coalescence sequence involving these lineages is *{* (A, B), (AB, C), (ABC, D), (ABCD, E) *}*, where A and B coalesce first, then C coalesces with the resulting AB lineage, then D coalesces with the resulting ABC lineage, and finally E coalesces with the resulting ABCD lineage.

Coalescence sequences involving disjoint sets of lineages can be combined into a single coalescence sequence that contains all the coalescences from both sequences, a procedure termed “interweaving” [e.g. Rosenberg, 2003]. The same set of coalescence sequences can be interwoven in different ways to form different interwoven coalescence sequences (Figure 3).

**Figure 3:**
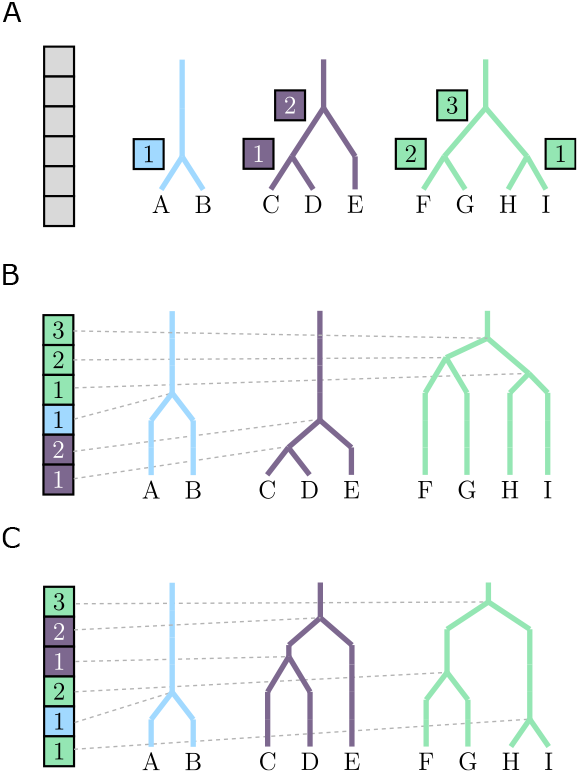
Interweaving of coalescence sequences. (A) Three coalescence sequences. The sequences are represented in three colors. Within a sequence, coalescences occur in a specified order, indicated by numbers within colors. Each of the six coalescences must occur in the interwoven sequence, represented by the gray blocks. Hence, each coalescence must be mapped to one of the gray blocks, with order increasing from bottom to top for each sequence. (B,C) Two different ways to interweave the sequences from (A).

### 2.6 Joint monophyly

Given a subtree *ℱ*_*x*_ of *ℱ*, defined as the node *x* and all of its descendants, we define the joint monophyly event *E*_*x*_, representing the event that joint monophyly occurs for gene lineages in the subtree *ℱ*_*x*_. For joint monophyly to be achieved, each coalescence in *ℱ*_*x*_ must be in one of four mutually exclusive classes:

1. The coalescence is between two lineages that have the same label (an intralabel coalescence), and neither label is mixed.
2. The coalescence is between two lineages with different labels (an interlabel coalescence), neither label is mixed, and both labels have only one existing lineage at the time of the coalescence.
3. The coalescence is between two lineages with different labels, exactly one of which is mixed, and the other label has only one existing lineage at the time of the coalescence.
4. The coalescence is between two mixed lineages.

### 2.7 Combinatorial functions

We use several combinatorial functions in our calculation. First, *g*_*i,j*_(*T*) is the probability that *i* lineages coalesce to *j* lineages in time *T*, from Eqn. 6.1 of Tavaré [1984]:

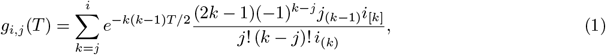

where *a*_(*k*)_ = *a*(*a* + 1) *…*(*a* + *k −* 1) and *a*_[*k*]_ = *a*(*a −* 1) *…*(*a − k* + 1) for *k* ≥ 1, and *a*_(0)_ = *a*_[0]_ = 1. This function is nonzero when *i* ≥ *j* ≥ 1 and *T* ≥ 0. We define *g*_0,0_(*T*) = 1, and we write *g*_*i*,1_(∞) for lim_*T* →∞_ *g*_*i*,1_(*T*).

Second, the number of coalescence sequences that reduce *n* lineages to *k* lineages is

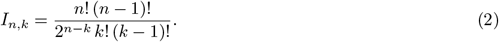

This function, from Eqn. 4 of Rosenberg [2003], is nonzero for *n* ≥*k*≥ 1, and we define *I*_0,0_ = 1.

Third, the multinomial coefficient

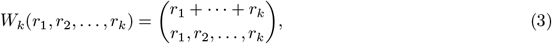

from Mehta and Rosenberg [2019], is the number of ways that *k* coalescence sequences of lengths *r*_1_, *r*_2_, …, *r*_*k*_ coalescent events can be ordered, or interwoven, to create an encompassing coalescence sequence that contains them all as subsequences. This function is defined for *r*_*i*_ ≥ 0, *i* = 1, 2, …, *k*.

Finally, *Z*(*s*_1_, *s*_2_, …, *s*_*k*_) is the probability that in a single population in which *k* groups are present, *k* groups of *s*_1_, *s*_2_, …, *s*_*k*_ gene lineages coalesce to a single lineage while preserving joint monophyly of each of the *k* groups. This function is reported by Theorem 5.1 of Zhu et al. [2011], as follows.

Suppose that *A*_1_, *A*_2_, …, *A*_*k*_ represent sets of lineages for groups 1, 2, …, *k*, respectively. Under joint monophyly of groups 1, 2, …, *k*, each group *i* possesses a single lineage *a*_*i*_ ancestral to all lineages in *A*_*i*_. The lineages *a*_*i*_ possess some labeled topology *T*_*k*_ from the set of all possible labeled topologies *rb*(*k*) (“rb” for rooted binary trees). We compute the probability that the *k* groups are jointly monophyletic and that their associated single-lineage ancestors possess labeled topology *T*_*k*_, and then sum over all possible *T*_*k*_ to obtain the total probability of joint monophyly of the *k* groups.

Let 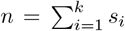 be the total number of lineages across all groups. Let *ℱ* (*T*_*k*_) be the set of internal nodes of *T*_*k*_. For an internal node *v* ∈ *ℱ* (*T*_*k*_), let *I*_*v*_(*A*_*i*_) denote the indicator function that lineage *a*_*i*_ is a descendant of *v* in *T*_*k*_. The joint probability of joint monophyly of the *k* groups and labeled topology *T*_*k*_ is

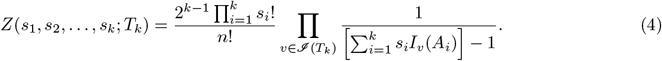

Summing over all (2*n −* 2)!*/*[2^*n−*1^(*n −* 1)!] possible *T*_k_ in *rb*(*k*), the total probability of joint monophyly is

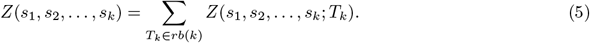

Our notation occasionally leads to values of 0 for some of the arguments of the function in Eqn. 5. In those cases, the quantity is properly computed by dropping those arguments.

## 3 Mathematical results

For species tree node *x*, we can compute the probability of the joint monophyly event *E*_*x*_ and a particular output state 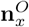 by recursive decomposition as follows:

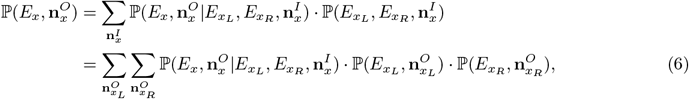

where *x*_*L*_ and *x*_*R*_ are the daughter nodes of *x*, and the second step is due to independence of these nodes and the fact that 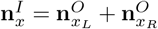. Taking *x* to be the species tree root, 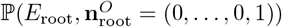 is the joint monophyly probability for the entire gene genealogy.

To compute ℙ (*E*_root_), we use a pruning algorithm—a familiar approach in phylogenetics in general [Felsenstein, 2004, p. 253]. First calculating the probability 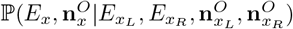—the probability of obtaining the joint monophyly event *E*_*x*_ and an output state 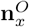 given the events 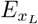 and 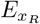 and their corresponding output states 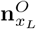 and 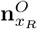 —we can apply this probability to the root of the tree and then proceed recursively to the leaves, whose inputs are known, ending the recursion. Given 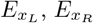, and their output states, the probability of event *E*_*x*_ and its output state is the probability that all coalescences that occur in the branch *x* satisfy joint monophyly and result in the specified output state. We can compute this probability by specifying an input state, computing the probability that joint monophyly is preserved on branch *x* given the input state and output state, and summing over all possible input states for branch *x*.

The probability that joint monophyly is preserved in a branch with a specified input state and output state requires computation of two quantities: (i) the probability that the correct number of coalescences occurs to convert the input state into the output state, and (ii) among coalescence sequences with the correct number of coalescences, the fraction that satisfy joint monophyly.

For (i), the probability that the correct number of coalescences occurs is 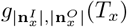 (see Section 2.7). For (ii), to count the coalescence sequences, the calculation is more involved. It is useful to first classify the *k* input labels into two categories: surviving and lost.

### 3.1 Surviving labels and lost labels

Consider a branch *x* with input state 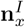 and output state 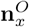. Consider a label *i* ∈ {1, 2, …, *k*}. The number of lineages of label *i* in the input state is denoted 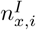 and its number of lineages in the output state is denoted 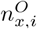. The total number of lineages in an input or output state is denoted 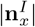 or 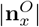, respectively.

Suppose 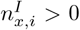. Two possibilities then exist for label 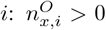 or 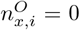. If 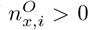, then label *I* is said to be a *surviving label* on branch *x*. To preserve joint monophyly on branch *x*, lineages of surviving label *i* are permitted to undergo intralabel coalescences on the branch, but not interlabel coalescences.

If 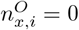, then label *i* is said to be a *lost label* on branch *x*. To preserve joint monophyly on branch *x*, lineages of lost label *i* undergo intralabel coalescences until only one lineage of label *i* remains. This final lineage undergoes an interlabel coalescence.

In the branch represented in Figure 2B, label 1 survives, whereas label 2 is lost.

### 3.2 Number of coalescence sequences for each surviving label

Our general approach for counting permissible coalescence sequences within a branch is to split the coalescences within the branch into multiple subsequences that we know how to count, and to then interweave those subsequences together. First, we consider sequences involving surviving labels. Under joint monophyly, each lineage in a surviving label must coalesce only with other lineages that possess that same label. Thus, the set of all input lineages of a particular surviving label *i*, the coalescences of those lineages, and the output lineages of label *i* can be used to define a coalescence subsequence for label *i*. The number of distinct coalescence subsequences for surviving label *i* is the number of ways that the 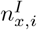 input lineages of label *i* can coalesce to the correct number of output lineages of label *i*, or 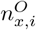. This number of subsequences is 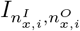 (Eqn. 2). We compute this quantity for each surviving label.

### 3.3 Enumerating partitions containing lost labels and mixed lineages

We next count coalescence subsequences that involve lost labels and mixed lineages. Unlike for surviving lineages, because a lost label must undergo an interlabel coalescence, coalescence subsequences involving lost labels only produce output mixed lineages. Hence, each output mixed lineage must result from a coalescence subsequence (i) involving at least two mixed lineages and no lost labels, (ii) involving at least two lost labels and no mixed lineages, or (iii) involving at least one lost label and at least one mixed lineage.

To account for every possible coalescence subsequence in one of these three categories, we must assign each output mixed lineage to an element of a partition of the set of lost labels and input mixed lineages.

Thus, we partition the *input* lineages, assigning to each element of the partition a single *output* mixed lineage. A coalescence subsequence exists for each element of the partition. We count the number of distinct types of lineages, among the input lineages with lost labels and the input mixed lineages. This quantity equals ℓ+ *m*_*I*_ : *ℓ* input lost labels and *m*_*I*_ individual input mixed lineages. The number of elements of the partition of output lineages is *m*_*O*_: one element for each of the *m*_*O*_ individual output mixed lineages. Thus, we are partitioning *ℓ* + *m*_*I*_ labeled elements into *m*_*O*_ nonzero categories. In particular, these partitions are the ways to place *ℓ* + *m*_*I*_ labeled balls into *m*_*O*_ unlabeled boxes, such that each box contains at least one ball [Loehr, 2017]. The number of these partitions are Stirling numbers of the second kind, *S*_2_(*ℓ* + *m*_*I*_, *m*_0_). An algorithm for producing these partitions is presented in Knuth [2011].

However, two additional conditions must be met.

1. *m*_*O*_ ≤ *m*_*I*_ + ⌊ℓ ⌋/2.
2. No element of the partition can consist solely of a single one of the *ℓ* lost labels.

In the first condition, the number of output mixed lineages is bounded above by the number of input mixed lineages plus the maximal number of additional mixed lineages that can be produced by coalescences involving the lost labels. Lost labels whose coalescences involve the *m*_*I*_ input mixed lineages do not generate additional output mixed lineages; however, lost labels whose coalescences involve other lost labels do generate additional output mixed lineages. The maximal number of output mixed lineages that can be generated in this way is ⌊ *ℓ ⌋ /*2, if the maximal number of pairs of lost labels coalesce with each other.

The second condition codifies the requirement that no output mixed lineage is generated purely by coalescences within a single lost label. Each lost label must coalesce with others or with mixed lineages.

Once all possible partitions of the *ℓ* + *m*_*I*_ labeled elements into *m*_*O*_ unlabeled nonempty sets are enumerated, we filter these partitions by the conditions 1 and 2, retaining only those partitions that satisfy both criteria. We define these partitions to be “permissible partitions.” For each partition retained, we next describe the enumeration of the coalescence subsequences associated with an element of the partition.

### 3.4 The number of coalescence subsequences for each element of a partition of the set of lost labels and mixed lineages

Denote by *𝒫* the set of permissible partitions of the set *L* ∪ *M*_*I*_, where *L* is the set of *ℓ* lost labels and *M*_*I*_ is the set of *m*_*I*_ input mixed lineages. Let *U* = *K \ L*, where *K* = *{*1, 2, …, *k}*. Let *P* be a partition in *𝒫*.

Consider an element *p* of *P*. This element is associated with a set *L*_*p*_ ⊂ *L* of lost labels and a set *M*_*p*_ of *m*_*p*_ input mixed lineages. *L*_*p*_ or *M*_*p*_ is possibly empty, but they cannot both be empty. Element *p* corresponds to a coalescence subsequence that starts with 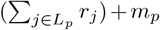 lineages and ends with a single mixed lineage, where *r*_*j*_ is the number of input lineages of (lost) label *j*.

Following Section 2.7, the number of subsequences that coalesce 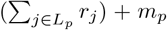 lineages to a single lineage is 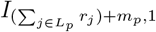 (Eqn. 2). The fraction of these subsequences that satisfy joint monophyly is *Z*(**v**_*p*_), where *Z* is the probability of joint monophyly of an arbitrary number of groups in a single population (Eqn. 5). The argument **v**_*p*_ is constructed as a vector of length *k* + *m*_*p*_. For elements *i* from 1 to *k, v*_*i*_ = *r*_*i*_ if *i* ∉ *L*_*p*_ and *v*_*i*_ = 0 if *i* ∈*/ L*_*p*_. The last *m*_*p*_ elements all equal 1. For example, consider a 7-species tree. If partition *p* contains lost labels 1 and 6 and three input mixed lineages, then **v**_*p*_ = (*r*_1_, 0, 0, 0, 0, *r*_6_, 0, 1, 1, 1).

Combining the number of subsequences that start from the input lineages in *p* and coalesce to a single lineage with the fraction of those subsequences that satisfy joint monophyly gives the total number of subsequences that both have the correct number of coalescences and that satisfy joint monophyly:

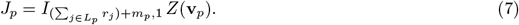

### 3.5 The number of coalescence sequences associated with a set of surviving labels and a partition of the set of lost labels and mixed lineages

Each of the |*U* | surviving labels and each element *p* of partition *P* creates a coalescence subsequence that must be interwoven with the other such subsequences. There are |*U* | + |*P* | such subsequences. For 1 ≤ *i* ≤ |*U* |, the number of coalescences is *s*_*i*_ *− r*_*i*_, noting that *U*_*i*_ is the *i*th surviving label (enumerated in arbitrary order) and abbreviating 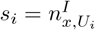 and 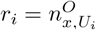 for convenience.

For each *i* with |*U*| + 1 ≤*i* ≤ |*U*| + |*P*|, the number of coalescences is |*P*_*i−*|*U*|_ |*−* 1coalescences, where *P*_*j*_ is the *j*th element of *P* (again enumerated in arbitrary order). Hence, the number of ways to interweave the |*U* | + |*P* | coalescence subsequences is (from Eqn. 3)

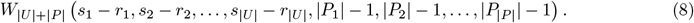

Multiplying the number of ways of interweaving the coalescence subsequences by the product of the numbers of ways of constructing the various subsequences, the total number of sequences that satisfy joint monophyly for a given input state and output state is

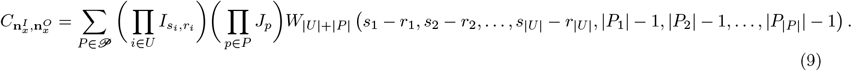

The product over elements of *U* is the number of coalescence sequences involving surviving labels. The product over elements of *P* is the number of coalescence sequences for a particular partition of lost labels and mixed lineages, and the sum over all *P* accounts for all possible partitions in𝒫.

If there are no surviving labels, then the product over elements of *U* is trivial, equal to 1. If all labels are surviving labels, then trivially, only a single partition in *P* ∈ *𝒫* is possible. We omit the sum over this partition *P*, and note that *J*_*p*_ = 1 trivially for the single element *p* of this trivial partition *P*. Eqn. 9 becomes

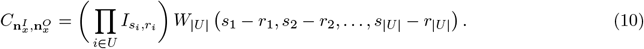

### 3.6 Completing the computation

The total number of coalescence sequences in a branch given an input state and an output state is 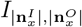 (Eqn. 2). The number that satisfy joint monophyly is 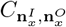, following Eqn. 9. From Eqn. 1, the probability of obtaining a particular number of coalescences in a branch of length *T*_*x*_ is 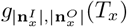.

We conclude that in Eqn. 6 for the probability of joint monophyly in branch *x* together with an output state, the recursive step that computes the conditional probability of joint monophyly and the output state given that joint monophyly is maintained in the daughter branches *x*_*L*_ and *x*_*R*_ and given the input state is

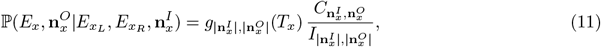

where 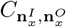 is from Eqn. 9 and 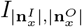 is from Eqn. 2. This result, applied recursively starting from *x* = root with 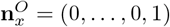, yields the probability of joint monophyly over all species 1, 2, … *k*.

### 3.7 Deriving previous results

We can use Eqn. 11 to derive previously-known results on the probability of joint monophyly under the multispecies coalescent. In this section, we proceed through several special cases.

#### 3.7.1 k groups in one population

This case has only one branch *x*, corresponding to the single population; *x* has no daughter nodes. There is only one possible input state into 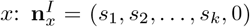, where *s*_*i*_ is the sample size of group *i*. The output state is 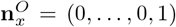, with size 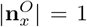. Branch *T*_*x*_ has infinite length. The summation in Eqn. 6 is trivial, and applying Eqn. 11, we obtain

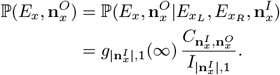

The labels 1, 2, …, *k* are all lost, and there is only one output mixed lineage *m*_1_. Hence, the set of partitions *𝒫* of lost labels and input mixed lineages into output mixed lineages consists of a single partition *P* = *{p}*, with single element *p* = *{*1, 2, …, *k}* → *m*_1_. Thus, when we use Eqn. 9, we obtain 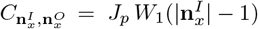. Noting that *g*_*i*,1_(∞) = 1, and from Eqn. 3, 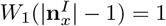, we find

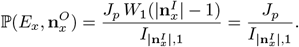

Using our notation from Section 3.4, the partition vector is **v**_*p*_ = (*s*_1_, *s*_2_, …, *s*_*k*_). We use Eqn. 7 to obtain

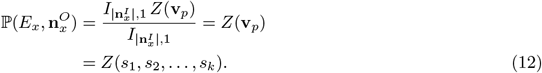

Note that *Z*(*s*_1_, *s*_2_, …, *s*_*k*_) is exactly the quantity in Eqn. 5, and we recover the result from Zhu et al. [2011].

#### 3.7.2 General term for a leaf node

Next, we consider a series of cases in which monophyletic groups correspond to the lineages of specific species (Table 1). A leaf node has exactly one input label *i* and exactly one surviving label *i*, and it has no other types of label. The input state is 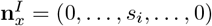, and the output state is 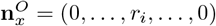.

**Table 1:** Analytical results for the probability of joint monophyly for arbitrary sample sizes. Eqn. 11 in Section 3.6 provides a general calculation from which we recover the other results listed. Other cases with small numbers of populations and monophyletic groups appear in Table 1 in Mehta and Rosenberg [2019].

| Study | Number of Populations | Number of Monophyletic Groups | Section |
| --- | --- | --- | --- |
| Zhu et al. [2011] | 1 | Arbitrary | 3.7.1 |
| Rosenberg [2003] | 2 | 2 | 3.7.3 |
| Mehta and Rosenberg [2019] | 3 | 3 | 3.7.4 |
| Mehta et al. [2016] | Arbitrary | 2 | 3.7.5 |
| This paper | Arbitrary | Arbitrary | 3.6 |

Thus, for a leaf node, using Eqn. 8, the partition set *𝒫* is trivial, producing Eqn. 10. The set of surviving lineages is *U* = *{i}*. Using Eqn. 10 along with Eqn. 3, we obtain

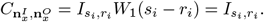

A leaf node has no daughter nodes, and the input state is therefore known; trivially, Eqn. 6 has a single term. Using Eqn. 11, we have

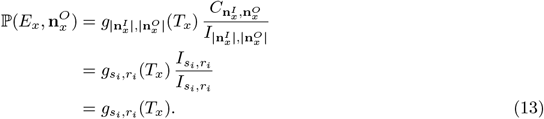

Thus, the general computation in Eqn. 11 reduces to Eqn. 1, the expression describing the probability that *s*_*i*_ lineages coalesce to *r*_*i*_ lineages in time *T*_*x*_.

#### 3.7.3 Two species in a two-species tree

In a two-species tree, let *s*_1_ and *s*_2_ be the initial sample sizes of species 1 and 2, respectively, and let *r*_1_ ≤ *s*_1_ and *r*_2_ ≤ *s*_2_ be the numbers of lineages of species 1 and 2 that enter the root node. There are three species tree nodes: the root *x*, leaf *x*_1_ for species 1, and leaf *x*_2_ for species 2. The input and output states are 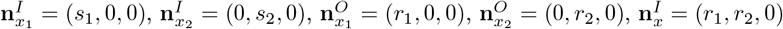, and 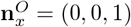.

For leaf *x*_1_, label 1 survives and there are no other label types. For leaf *x*_2_, label 2 survives and there are no other label types. For the root, both species labels are lost, and there is only one output mixed lineage *m*_1_. Hence, there is only one partition *P* = {{1, 2 } →*m*_1_}.

Because *x*_1_ and *x*_2_ are leaves, from Eqn. 13, 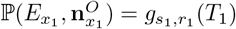 and 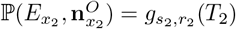. From Eqn. 12, for a particular *r*_1_ and *r*_2_, we have 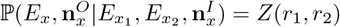. Substituting into Eqn. 6,

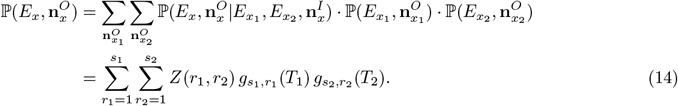

It remains to obtain *Z*(*r*_1_, *r*_2_). First, note that there is only one possible labeled topology *T*_2_ for the two ancestral lineages of the two groups *A*_1_ and *A*_2_, and this topology has a single internal node *v* of which both *A*_1_ and *A*_2_ are descendants. So, for *k* = 2, we have by Eqns. 4 and 5,

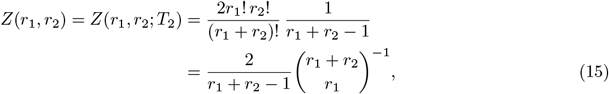

which matches Lemma 4.3 in Zhu et al. [2011], Eqn. 6 in Brown [1994], and Eqn. 9 in Rosenberg [2003]. Substituting Eqn. 15 into Eqn. 14, we have

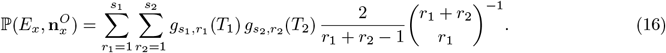

We therefore obtain Eqn. 14 from Rosenberg [2003]: the probability of reciprocal monophyly of two species in a two-species tree.

#### 3.7.4 3 species in a 3-species tree

In this section, we recapitulate the probability of joint monophyly for 3 species in a 3-species tree, as provided in Eqn. 5 in Mehta and Rosenberg [2019]. It suffices to describe the reduction of our Eqn. 11 to Eqns. 6 and 9 in Mehta and Rosenberg [2019], giving the conditional probability of joint monophyly within the internal node *I* of *ℱ* given a particular input state 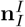 and output state 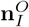, and the conditional probability of joint monophyly in the species tree root *R* given a particular input state 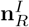.

We label the three leaves A, B, and C, and we call the single internal node *I* (ancestral to species A and B). The root node is *R*. Thus, we can specify branch input and output states:

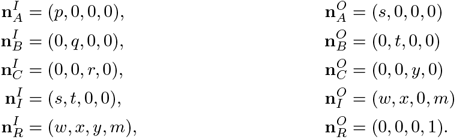

Eqns. 6 and 9 in Mehta and Rosenberg [2019] are special cases of a term in Eqn. 3 from Mehta and Rosenberg [2019], which corresponds to our Eqn. 11. Comparing Eqn. 11 to Eqn. 3 from Mehta and Rosenberg [2019] indicates that to obtain Eqn. 6 of Mehta and Rosenberg [2019], we must show that the quantity *K*_*I*_ from Mehta and Rosenberg [2019] satisfies

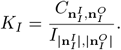

To obtain Eqn. 9 from Mehta and Rosenberg [2019], we must show that with **v**_*p*_ = (*w, x, y, m*), the quantity *K*_root_ from Mehta and Rosenberg [2019] satisfies

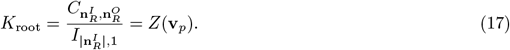

First, we consider internal node *I*. The nontrivial cases of Eqn. 6 from Mehta and Rosenberg [2019] are:

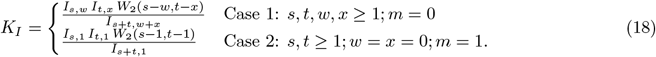

Eqn. 18 concerns the internal node *I* of a three-species tree, a node that has input lineages from the two species it subtends. Case 1 in Eqn. 18 occurs when both species labels are surviving labels, as the two quantities that represent the numbers of output lineages from the two input species, *w* and *x*, are both greater than or equal to 1. In the language of our analysis, the set of surviving labels is *U* = *{*1, 2*}*. There are no output mixed lineages (*m* = 0) and there is no need to consider a set of partitions *𝒫* of the set of lost labels and mixed lineages.

We use Eqn. 10 to obtain

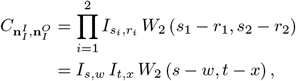

and so we have:

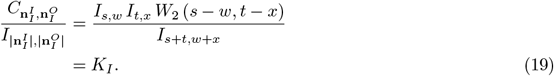

Case 2 in Eqn. 18 occurs when both species labels are lost labels, as the two quantities that represent the number of output lineages from the two input species, *w* and *x*, are both 0. There is one output lineage, a mixed lineage (*m* = 1). There is only one possible partition of input labels *{*1, 2*}* over the single mixed lineage *m*_*I*_ : *P* = *{p}*, with *p* = *{*1, 2*} → m*_*I*_. We use Eqn. 9 and Eqn. 7 to obtain:

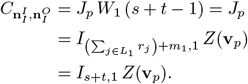

The vector **v**_*p*_ is (*s, t*, 0, 0). Note that *Z*(*s, t*, 0, 0) = *Z*(*s, t*), so we can use Eqns. 15, 2, and 3 to obtain

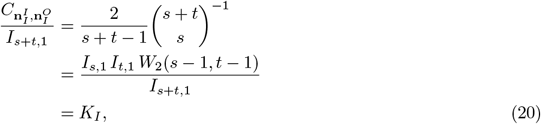

where the last step comes from Eqn. 18. Hence, we have 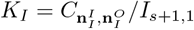, as desired.

It remains to show that our result accords with the two nontrivial cases of Eqn. 9 from Mehta and Rosenberg [2019]. These cases are:

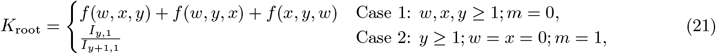

where

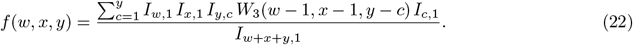

Starting from our Eqn. 17, we must calculate *Z*(**v**_*p*_) for each of these two cases and show that it equals *K*_root_ from Eqn. 21. Case 1 of Eqn. 21 occurs when there are input lineages from three species (*w, x, y ≥* 1) and no input mixed lineages (*m* = 0). Thus, **v**_*p*_ = (*w, x, y*, 0). We note that *Z*(*w, x, y*, 0) = *Z*(*w, x, y*). From an unlabeled example in Zhu et al. [2011] immediately following the proof of their Theorem 5.1, we have

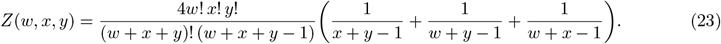

Substituting Eqns. 2 and 3 into Eqn. 22 and simplifying, we have

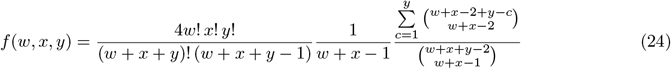

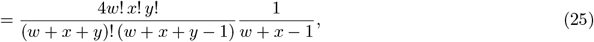

where the step from Eqn. 24 to Eqn. 25 uses the binomial identity (Eqn. 1 in Section 0.151 from Gradshteyn and Ryzhik [2014])

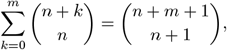

with *y − c* in place of *k, y −* 1 in place of *m*, and *w* + *x −* 2 in place of *n*.

Now, from Eqns. 23 and 25, we have

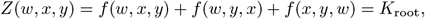

as required.

Case 2 of Eqn. 21 occurs when there are input lineages from one species (*y ≥* 1, *w* = *x* = 0) and one input mixed lineage (*m* = 1), **v**_*p*_ = (0, 0, *y*, 1). We note that *Z*(0, 0, *y*, 1) = *Z*(*y*, 1), and use Eqn. 15 to obtain:

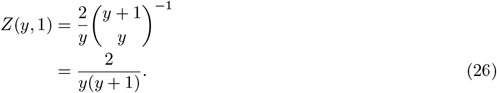

Using Eqns. 2 and 26, we have *Z*(*y*, 1) = *I*_*y*,1_*/I*_*y*+1,1_ = *K*_root_, as required.

Having demonstrated that our joint monophyly calculation recovers the combinatorial terms *K*_*I*_ and *K*_root_, we have therefore recapitulated the joint monophyly probability for three species, as obtained by Mehta and Rosenberg [2019].

### 3.7.5 2 groups in a *k*-species tree

Here we recapitulate the probability of joint monophyly for 2 groups in a *k*-species tree, as shown in Eqn. 5 from Mehta et al. [2016]. It suffices to describe the reduction of our Eqn. 11 to Eqn. 4 from Mehta et al. [2016], describing the conditional probability of monophyly within the a node *x* of *𝒯* given a particular input state 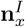 and output state 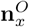. More precisely, we must equate our Eqn. 11 to the scenario of joint monophyly in Eqn. 4 of Mehta et al. [2016], obtained by substituting their Eqn. 5 for Case 2 in their Eqn. 4.

Let the input state for node *x* be 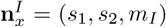, and let the output state be 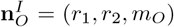. We assume (as is necessary to achieve joint monophyly) that the input lineages from groups 1 and 2 include all lineages from those groups; that is, species tree node *x* is ancestral to all lineages that belong to groups 1 and 2. Following the labeling of cases in Mehta et al. [2016], the nontrivial cases of Eqn. 4 from Mehta et al. [2016] in the setting of joint monophyly are:

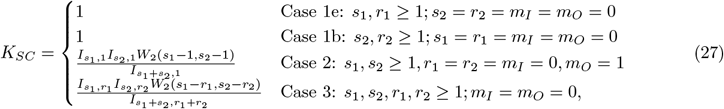

where Cases 1b and 1e in Eqn. 27 are labeled after their corresponding labels in Mehta et al. [2016].

Eqn. 4 in Mehta et al. [2016] is a special case of a term in Eqn. 3 from Mehta et al. [2016]. Comparing our Eqn. 11 to Eqn. 3 from Mehta et al. [2016], we find that to obtain Eqn. 4 of Mehta et al. [2016] as a special case of our Eqn. 11, we must show that the quantity *K*_*SC*_ from Mehta et al. [2016] satisfies

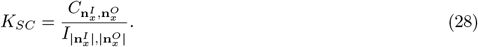

Cases 1e and 1b from Eqn. 27 occur when there is one surviving label and no other input lineages. We have *U* = *{i}* for *i* = 1, 2 and *𝒫* empty. We use Eqns. 10, 2, and 3 to obtain:

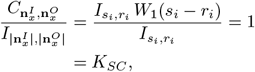

as required for demonstrating Eqn. 28.

Case 2 from Eqn. 27 occurs when there are two lost labels, no surviving labels, and one output mixed lineage *m*_*I*_. Thus, *U* is empty, and there is one partition *P* = {{1, 2} *→ m*_*I*_ }. We have already shown that Eqn. 11 produces Eqn. 20; directly applying the result from Eqn. 20 yields the result that Eqn. 28 requires. Case 3 from Eqn. 27 occurs when there are two surviving labels, no lost labels, and no input or output mixed lineages. Thus, *U* = {1, 2} and *𝒫* is empty. We have already shown that Eqn. 11 produces Eqn. 19; directly applying the result from Eqn. 19 yields the result required for Eqn. 28 to be satisfied.

We have therefore shown that our Eqn. 11 reduces to Eqn. 28, recapitulating the joint monophyly probability of two groups in an arbitrary species tree from Mehta et al. [2016].

### 3.8 Lower and upper bounds based on “strong” joint monophyly

The probability in Eqn. 11 involves many steps and is potentially time-consuming to calculate. We can therefore provide a simpler lower bound by introducing the idea of “strong” joint monophyly. We say that a set of lineages sampled from a species satisfies *strong joint monophyly* if the lineages coalesce to a single lineage in the branch associated with that species. The probability of strong joint monophyly can then be computed from the lengths of these branches, so that the rest of the tree structure does not matter.

The probability of strong joint monophyly is

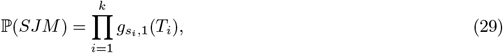

where *T*_1_, *T*_2_, …, *T*_*k*_ are the species tree branch lengths associated with species 1, 2, …, *k*.

This probability provides a lower bound on Eqn. 11 because it is only one of many ways that joint monophyly can be achieved; if *JM* denotes the event of joint monophyly, then ℙ (*JM*) *≥* ℙ (*SJM*). This lower bound avoids the pruning step and does not need to track lineage counts at species tree internal nodes, so that its calculation is faster than that of Eqn. 11. The lower bound is similar in spirit to an upper bound on the probability of gene-tree-species-tree concordance found by Pamilo and Nei [1988].

We can also observe that ℙ (*SJM*) enables an *upper* bound on ℙ (*JM*), a bound that holds for any species tree and any distribution of gene lineages across species. This bound is:

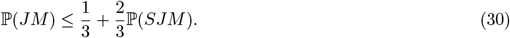

To prove Eqn. 30, first observe that if each species has exactly one lineage, then Eqn. 30 is an equality (i.e. 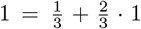). Thus, we can suppose that at least one species has at least two lineages, so that ℙ (*¬SJM*) *>* 0 and ℙ (*JM* |*¬SJM*) is well-defined. In this case, by the law of total probability,

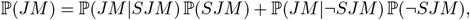

and because ℙ (*JM* |*SJM*) = 1, we obtain:

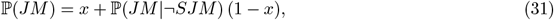

for *x* = ℙ (*SJM*). Next, we claim that:

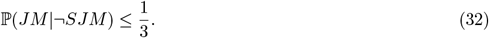

The justification of Eqn. 32 is as follows. The coalescent scenarios that comprise the event *¬SJM* are precisely those for which, for some tip species *s*, the (two or more) lineages associated with *s* do not coalesce to a single lineage within the pendant edge incident with *s*. However, joint monophyly (*JM*) requires that the ancestral lineages of *s* coalesce only among themselves (and not with other lineages) until they reach a single lineage along the path in *𝒯* back to its root. At some point on this path there will be just two ancestral lineages of *s*, along with *r ≥* 1 other ancestral lineages from other species, and the next coalescence will involve at least one of the two from *s*. The probability that the two ancestral lineages of *s* coalesce with each other (rather than one coalescing with one of the other *r* lineages present) is 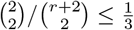 for all *r ≥* 1. Thus, 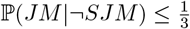 as claimed.

Combining Eqns. 31 and 32 gives Eqn. 30.

## 4 Numerical results

### 4.1 Continuous-time Markov Chain approach

Although the exact computation in Section 3.6 is instructive and presents new mathematical insight, using this result for computational purposes is inconvenient. To facilitate computation of the probability of joint monophyly, we provide a continuous-time Markov chain (CTMC, Grimmett and Stirzaker [2020]) formulation of this probability, described in Appendix A. When constructing this CTMC under the multispecies coalescent, we follow the approach of Hobolth et al. [2011]. Computing the probability of joint monophyly amounts to using the same recursive decomposition as in Eqn. 6, but the probabilities are computed by constructing transition matrices for each branch of the tree and using matrix exponentials to obtain the output probabilities given the input probabilities. We use the CTMC formulation in providing numerical results in this section. The approach is implemented in Monophyler [Mehta et al., 2016].

### 4.2 Effects of number of species, tree height, and sample size

We use example species trees to illustrate the effects of tree height and sample size on the probability of joint monophyly using an example. We consider a class of species trees that appears in in Figure 4. The trees range in size from two to six species, and they are constructed so that the tree height is evenly divided along the branches of the longest topological path length from root to leaf.

**Figure 4:**
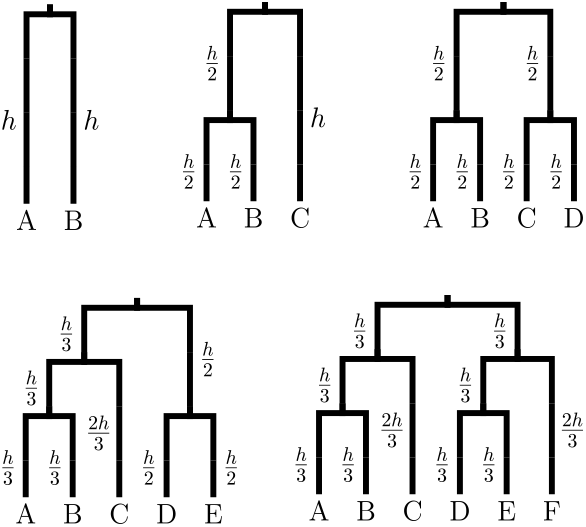
Trees used to explore the effects of tree height and sample size on the probability of joint monophyly.

Using each tree in Figure 4, we compute the probability of joint monophyly with Eqn. 11. We modulate the tree height *h* from 0 to 10 coalescent time units at intervals of 0.2. The number of samples in each leaf ranges from 2 to 10, incremented by 1, with each leaf having the same sample size.

Figure 5 displays the effect of number of species, tree height, and sample size on the probability of joint monophyly for all trees in Figure 4. As the number of species increases from 2 to 6, the number of separate groups that must be monophyletic in order to produce joint monophyly increases. Hence, the joint monophyly probability decreases at fixed values for the tree height and sample size.

**Figure 5:**
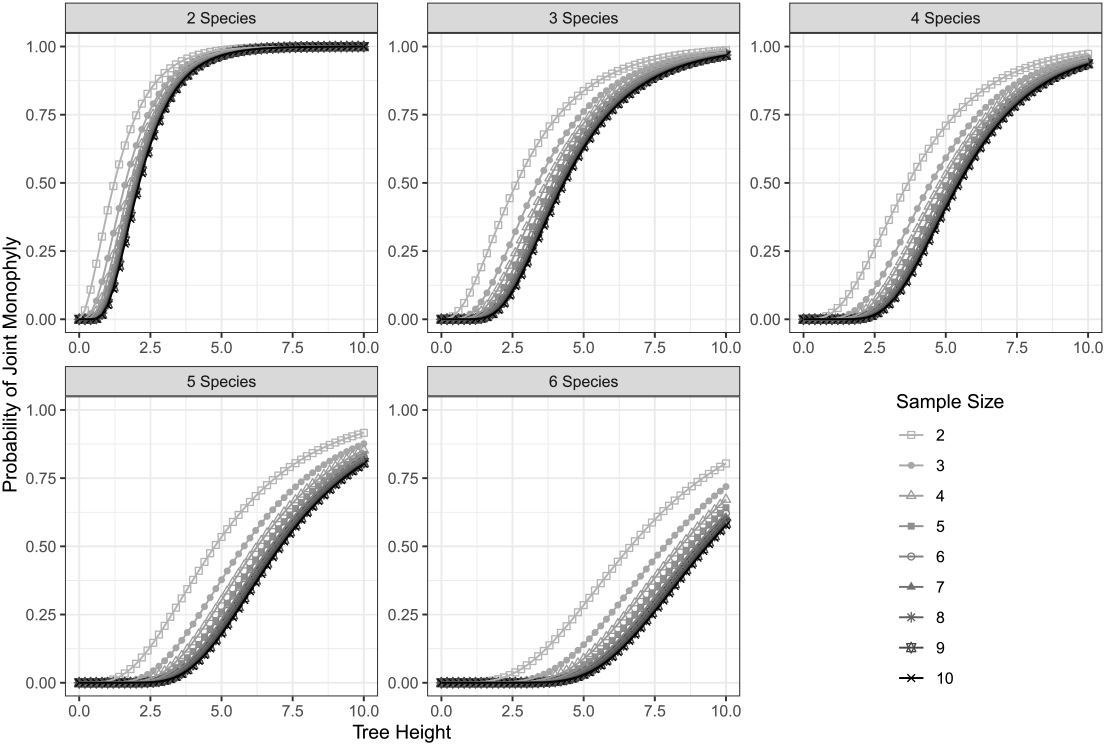
Joint monophyly probabilities for various numbers of species, tree heights, and sample sizes. Probabilities are obtained using Eqn. 11, with the same sample size assigned to each species. Each panel is labeled by the number of species.

With increasing tree height and fixed sample size, lineages have more time during which they can coalesce within the species from which they have been sampled, and the joint monophyly probability increases with increasing tree height. As the sample size increases at a fixed tree height, the number of lineages that must monophyletically coalesce increases, but no additional time is available for these coalescences; hence, the joint monophyly probability decreases with increasing sample size.

An alternative perspective on the joint monophyly probabilities in Figure 5 examines, for a fixed cutoff value representing a level of statistical significance, a fixed number of species, and a fixed tree height, the minimum sample size required for achieving a joint monophyly probability that lies below the cutoff. In other words, we calculate the minimum sample size required for an observation of joint monophyly to be improbable at a specified significance level under a specified model. Such a computation can assist in understanding the extent to which an observation of monophyly can be regarded as surprising and in designing samples such that a desired level of “surprise” is achieved if joint monophyly is observed [Rosenberg, 2007].

Figure 6 plots these minimum sample sizes. They decrease as the cutoff value is increased. In accord with the decrease in joint monophyly probabilities that occurs with an increasing number of species, for fixed tree height, the minimal sample size required for achieving a joint monophyly probability below a specified cutoff decreases with an increasing number of species. The minimal sample size increases with increasing tree height; as tree height grows, joint monophyly is probable even for large samples, so that very large samples might be required for a joint monophyly observation to be surprising. In most scenarios plotted, samples of 6 to 8 per species suffice to produce probabilities below cutoff 0.001 over most of the domain for tree height.

**Figure 6:**
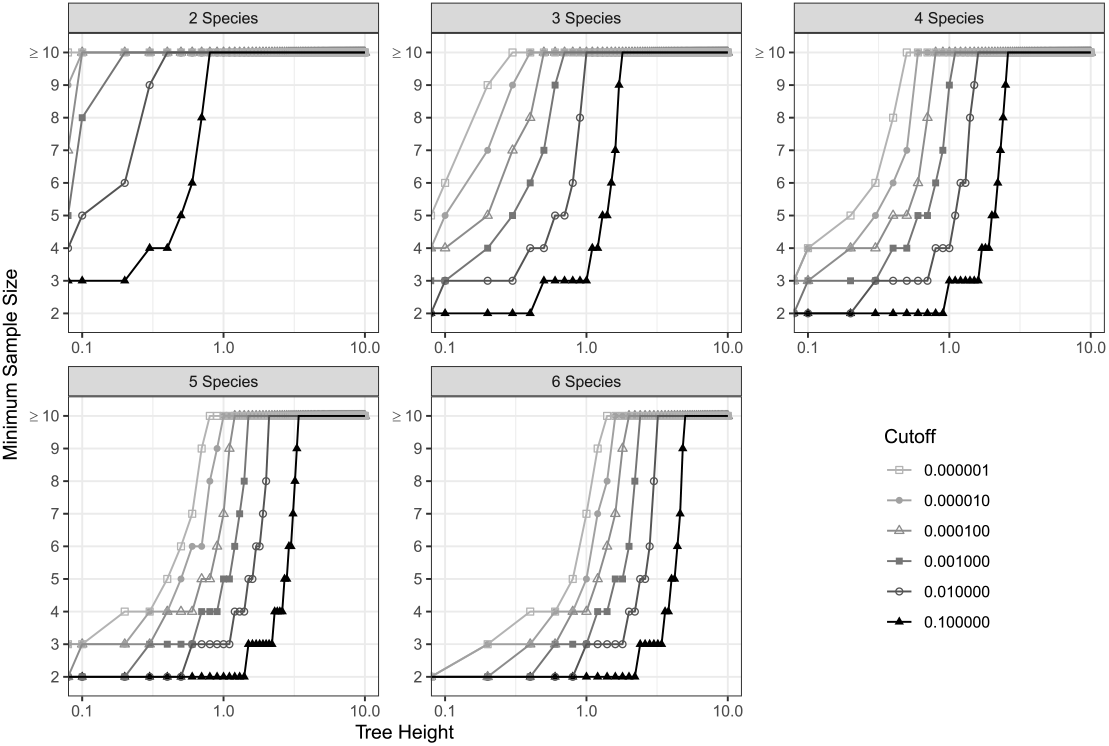
Minimum sample sizes for the probability of joint monophyly to decrease below a particular cutoff probability, for varying tree height and number of species. Panel title indicates number of species.

### 4.3 Strong joint monophyly

Figure 7 displays the probability of joint monophyly against the corresponding probability of strong joint monophyly from Eqn. 29. For each combination of a number of species, tree height, and sample size considered in Figure 5, the probability of strong joint monophyly is calculated, and a point is plotted that pairs the probability of strong joint monophyly with the probability of joint monophyly from Figure 5.

**Figure 7:**
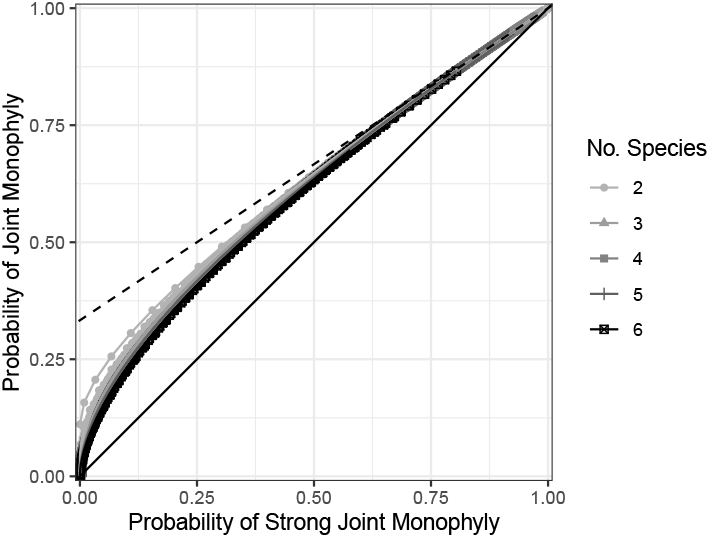
The probability of joint monophyly (Eqn. 11) in relation to the probability of strong joint monophyly (Eqn. 29). Strong joint monophyly provides a lower bound for joint monophyly. For each combination consisting of a number of species (2 to 6) and a sample size (2 to 10), a curve links points with increasing tree height (0 to 10 at intervals of 0.2). Parameter sets (number of species, tree height, sample size) follow Figure 4. The solid line indicates equality of the probabilities of joint monophyly and strong joint monophyly, and the dashed line indicates the upper bound on joint monophyly provided by Eqn. 30.

As strong joint monophyly is a stricter condition than joint monophyly, the probability of strong joint monophyly is necessarily less than or equal to the probability of joint monophyly (Section 3.8). Traversing the figure from left to right, or from bottom to top, the tree height increases. For large tree heights, joint monophyly is closely approximated by strong joint monophyly, as represented by the proximity of the curves plotted to the *y* = *x* line; the event of strong joint monophyly is the primary driver of joint monophyly. For smaller tree heights, the probability of strong joint monophyly is substantially lower than than the probability of joint monophyly, as configurations in which joint monophyly is achieved by coalescences that occur deeper in the species tree than the external branches are not improbable.

The plots show relatively little effect of the number of species on the relationship between joint monophyly and strong joint monophyly, or of the sample size. Thus, by curve-fitting, it would be possible to empirically transform the easily-computable SJM probability that approximates the joint monophyly probability.

For the case of 2 species and 2 lineages per species, using the 2-species tree in Figure 4, the probability of joint monophyly (JM) from Eqn. 16 is

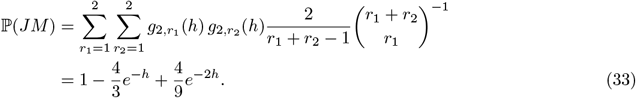

The probability of strong joint monophyly from Eqn. 29 is ℙ (*SJM*) = *g*_2,1_(*h*)^2^ = (1 *− e*^*−h*^)^2^. Solving this equation for *e*^*−h*^ and inserting the solution into Eqn. 33, the probability of joint monophyly (JM) in terms of the probability of strong joint monophyly (SJM) is

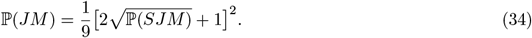

Eqn. 34 appears in Figure 7 as the curve corresponding to two species and sample size two, visible as the curve with the highest values of the JM probability for low values of the SJM probability.

## 5 Discussion

We have derived the general probability of joint monophyly in an arbitrary species tree—the probability that for each species in a *k*-species tree, the lineages of that species are monophyletic—under the multispecies coalescent. Using this result (Eqn. 11), we have obtained as special cases several previous results for the probability of joint monophyly: the cases of arbitrarily many groups of lineages in one species (Section 3.7.1), two lineage groups in two species (Section 3.7.3), three lineage groups in three species (Section 3.7.4), and two lineage groups in arbitrarily many species (Section 3.7.5). Previous results on the probability of joint monophyly were restricted to small numbers of groups (4 or fewer), small trees (4 species or fewer), or both. We were able to fully generalize these results by combining the recursive approach of Mehta et al. [2016] for general species trees and the combinatorial calculations of Zhu et al. [2011] for arbitrary numbers of groups.

Our calculation relies on a “pruning algorithm,” in which computations are performed recursively at each internal node of a species tree. Pruning algorithms have a long history in phylogenetics, tracing to early efforts to evaluate gene tree probabilities from molecular sequence data in maximum-likelihood phylogenetics [Felsenstein, 1981]. Recent algorithms have generalized the pruning approach to gene tree computations conditional on species trees [Efromovich and Kubatko, 2008, RoyChoudhury et al., 2008, Bryant et al., 2012, RoyChoudhury and Thompson, 2012, Stadler and Degnan, 2012, Wu, 2012, Mehta et al., 2016]. The pruning algorithm we have provided accounts for the intricate merging pattern of gene lineages that occurs when two species merge backward in time to their ancestral species.

Although pruning algorithms do lead to exact computations for various quantities of interest, they can suffer from the computational burden of tree traversal as the size of the species tree increases. In addition, though the pruning algorithm renders the tree traversal to be polynomial-time in the number of species, the computation time is not polynomial-time in the number of species or sample size, due to the effect on the most computationally complex part of the calculation: enumerating partitions and performing a calculation for each partition (Section 3.7.4). Our analysis includes the instructive formal computations that appear in Section 3 as well as a continuous-time-Markov-chain approach that is convenient for computation (Appendix A). Using the CTMC approach, we have seen that the joint monophyly calculation reproduces sensible patterns in the effects of model parameters on monophyly probabilities.

The new algorithmic approach will be useful in problems where joint monophyly of multiple groups is of interest—such as in phylogeographic and species delimination problems that have been examined in taxon groups including rotifers [Birky et al., 2005], birds [Cloutier et al., 2019], and snakes [Kubatko et al., 2011], among others. We have implemented the new algorithms in the software Monophyler [Mehta et al., 2016].

## A Calculating probabilities with a CTMC approach

### A.1 Mathematical approach

We now produce an alternative approach to calculating the probability of joint monophyly: a continuous-time Markov chain (CTMC) [Grimmett and Stirzaker, 2020]. We define a transition-rate matrix for each species tree branch and traverse the species tree from the leaves to the root. For each branch of the traversal, we use the probability of the input states and the transition-rate matrix to obtain the probability of the output states. The output states of two daughter branches combine to form input states of the parent branch.

Each branch of the species tree has its own Markov chain. For a particular branch *x*, we first must define a state space. Let *𝒯*_*x*_ be the subtree below and including branch *x*. For this appendix, we track lineage labels differently from Section 3. We no longer keep track of “lost,” “surviving,” or “mixed” labels. Instead, we classify the species labels *{*1, 2, …, *k}* by their numbers of extant lineages. A label *i* for one of the *k* species starts at a leaf with *s*_*i*_ lineages—the sample size of the species. If joint monophyly is preserved, then the *s*_*i*_ lineages eventually decrease to a single ancestral lineage. Once the single lineage is reached, the label and its single associated extant lineage become “free,” in that any coalescence involving this label no longer affects its contribution to joint monophyly. Coalescences of free lineages with other free lineages preserve joint monophyly, reducing the number of free lineages. In this formulation, “mixed” lineages are free.

The state space for a branch *x* therefore consists of a “failure” state F, which represents the situation where joint monophyly has been violated, and a set of vectors **v**_*x*_ that keep track of the list of lineage counts for the *k* labels. The *i*th element of **v**_*x*_, *v*_*x,i*_, is the number of *labels* with *i extant lineages*, with *v*_*x*,1_ counting the number of free lineages. For a branch *x*, the maximum number of lineages a label can have is the largest sample size of any species in *𝒯*_*x*_, as no label can gain lineages through coalescence. If *S*_*x*_ is the set of species in *𝒯*_*x*_, then the vectors in the state space for the chain for branch *x* have length 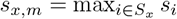.

State transitions in this process occur due to coalescence. Let us define

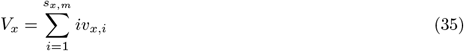

as the total number of lineages for state *v*_*x*_. For state **v**_*x*_, we have three possible transitions, corresponding to intralabel coalescences, interlabel coalescences that preserve joint monophyly, and interlabel coalescences that do not preserve joint monophyly.

1. An intralabel coalescence within a label of size *i >* 1 reduces the number of lineages of that label by 1. 1. *v*_*x,i*_ *→ v*_*x,i*_ *−* 1, and *v*_*x,i−*1_ *→ v*_*x,i−*1_ + 1. There are 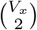 possible coalescences, and among those, 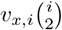 lead to this state transition. The conditional probability that a coalescence has this transition given that a coalescence occurs is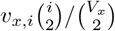.
2. An interlabel coalescence that preserves joint monophyly can only occur between free lineages. Thus, it reduces *v*_*x*,1_ *→ v*_*x*,1_ *−*1. The conditional probability that a coalescence has this transition is 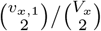.
3. Finally, any other coalescence is an interlabel coale scen ce tha t v iolates join t m ono phyly. Hence, *v*_*x*_ *→ F*. This transition has conditional probability 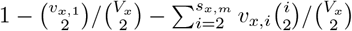.

These probabilities yield a transition matrix for transitions conditional on occurrence of a coalescence.

### A.2 Example transitions and transition rate matrices

Consider a branch with input lineages from four species—two with 1 lineage, one with 2 lineages, and one with 4 lineages—as well as a single input mixed lineage. There are three “free” lineages: those of species 1 and 2 and the mixed lineage. The input state is **v**_*x*_ = (3, 1, 0, 1). The total number of lineages present is *V*_*x*_ = 9 (Eqn. 35), so there are 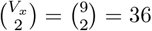 possible coalescences. The maximal sample size is *s*_*x,m*_ = 4.

Four types of coalescences are possible. An intralabel coalescence can occur in the species with 2 lineages, *i* = 2. This coalescence has probability

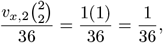

This transition converts a species with 2 lineages to one with 1 lineage, or (3, 1, 0, 1) *→* (4, 0, 0, 1).

An intralabel coalescence can also occur in the species with 4 lineages. In this case, *i* = 4, so that the transition probability is

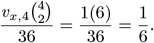

The species with 4 lineages transitions to one with 3 lineages. The state transition is (3, 1, 0, 1) → (3, 1, 1, 0). Interlabel coalescences can occur between free lineages (*i* = 1). This transition occurs with probability

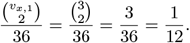

It reduces 3 free lineages to 2 free lineages, and the state transition is (3, 1, 0, 1) *→* (2, 1, 0, 1). Finally, any other coalescence leads to the failure state. Hence, (3, 1, 0, 1) *→ F* with probability

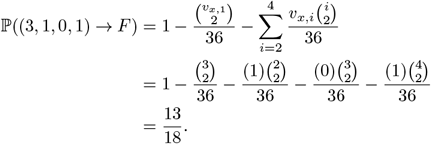

The full transition matrix for this species tree branch includes every state attainable by any number of coalescences beginning with state (3, 1, 0, 1). Figure 8 displays the state space for the branch, along with all possible transitions. The complete transition matrix can be obtained by using similar reasoning for all possible states and appears in Table 2.

**Figure 8:**
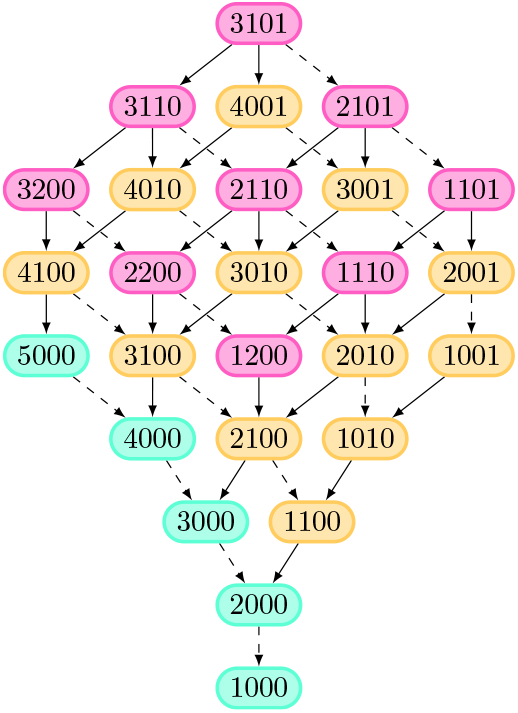
State space for the continuous-time Markov chain for the example branch in Section A.2. States are colored by the number of species for which joint monophyly is not yet determined (pink, two; yellow, one; green, none). Intraspecies transitions use a solid line; interspecies transitions use a dashed line. The Failure state is excluded; all states except those colored green can transition to the failure state.

**Table 2:**
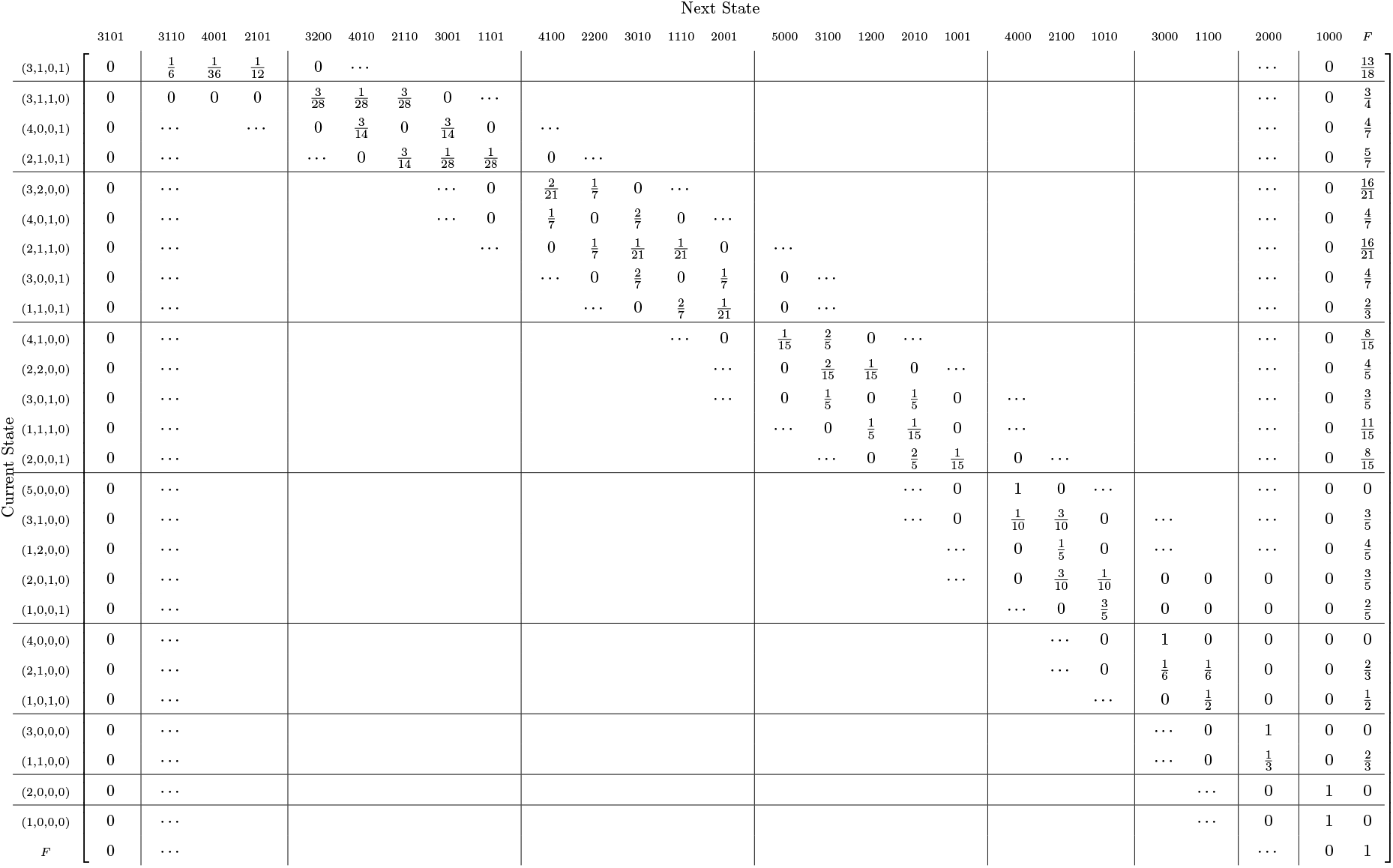
Transition matrix for the example continuous-time Markov chain in Section A.2.

To get the CTMC, the transition matrix for a branch must be converted into a transition rate matrix, or a Q matrix. We first scale the transition rates by noting that for a state **v**_*x*_, coalescences occur at rate *V*_*x*_. Next, we subtract each row sum from the associated diagonal entry of the matrix. Thus, to obtain our transition rate matrix, we must first multiply each row by the total number of possible coalescences for the state corresponding to that row, and then subtract the row sum from the diagonal entry. Therefore, the Q matrix for the branch follows the matrix in Table 3.

**Table 3:**
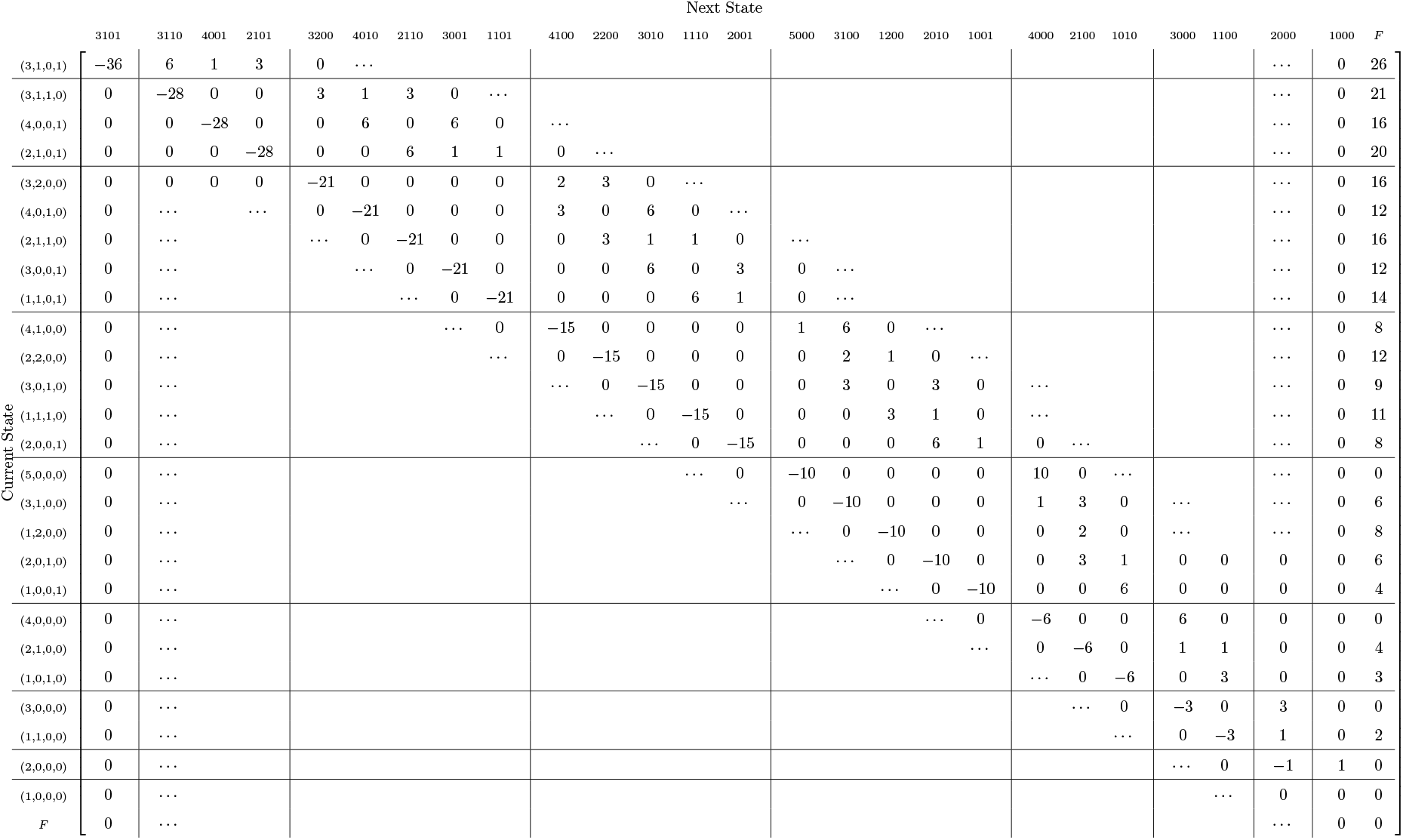
Transition rate matrix (*Q*_*x*_) for the example continuous-time Markov chain in Section A.2.

Given a vector of probabilities *p*_*x*_ that represents the input probability distribution over all possible states in a species branch *x* with length *T*_*x*_, the distribution of output states is [Grimmett and Stirzaker, 2020]

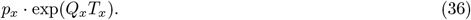

### A.3 Algorithm

The CTMC algorithm consists of two components. First, the species tree structure is created. Second, a recursive function is applied to the root node of the tree, and this function returns the probability result. The following pseudocode describes the creation of the tree structure:

read tree from Newick string;

read sample size information;

assign sample sizes to leaves of tree;

do recursive function `getnodeoutput` on root node of tree;

The tree structure is created within Python from a string in Newick format by using the function `Tree` in the package `ete3`. The user must specify both the Newick tree and the sample size information, which consists of two lists: one specifying the leaf names (the same names as in the Newick tree), and the other specifying the sample sizes of those leaves in the same order.

The recursive function `getnodeoutput` is described in the following psuedocode:

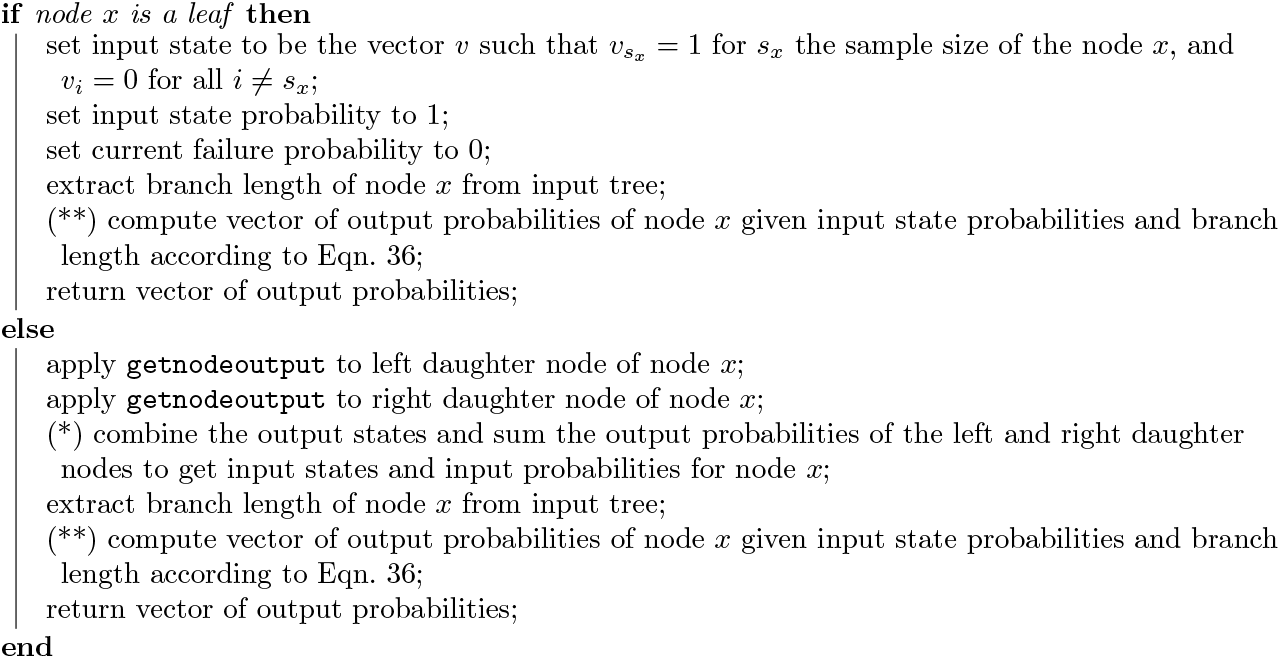

**Step (*): combining inputs**. Step (*), “combining” the output states and summing the output probabilities of the left and right daughter nodes *L* and *R*, respectively, of a node *x*, proceeds as follows.

We first note that there are no shared species labels between daughter nodes. Hence, all species labels with *i* extant lineages in the output of node *L* or node *R* also have *i* extant lineages in the input of node *x*. The number of species labels with *i* extant lineages as inputs of node *x, v*_*x,i*_, is *v*_*L,i*_ + *v*_*R,i*_ for each *i >* 1. Similarly, free lineages in the output states of nodes *L* and *R* remain free as input lineages to *x*, so *v*_*x*,1_ = *v*_*L*,1_ + *v*_*R*,1_. Thus, summing vectors, an input state **v**_*x*_ obtained from a pair of output states **v**_*L*_ and **v**_*R*_ is **v**_*x*_ = **v**_*L*_ + **v**_*R*_. The probability of the input state **v**_*x*_ obtained by summing output states **v**_*L*_ and **v**_*R*_ is the product of the probabilities of the output states **v**_*L*_ from node *L* and **v**_*R*_ from node *R*.

The set of possible input states to node *x* is obtained by considering all possible sums of an output state for daughter node *L* and daughter node *R*, using vector summation. The probability of a possible input state to *x* is the sum over all pairs of output states that result in that input state, where for each pair, the product of the probabilities of the two output states in that pair is summed.

This vector addition procedure omits the failure state *F*, which occurs as an input state of node *x* when it is an output state of *L, R*, or both *L* and *R*. If ℙ (*F*)_*L*_ and ℙ (*F*)_*R*_ are the output failure probabilities for *L* and *R*, respectively, then the input probability of failure is ℙ (*F*)_*L*_ [1 *−*ℙ (*F*)_*R*_] +[1 *−*ℙ (*F*)_*L*_] ℙ (*F*)_*R*_ + ℙ (*F*)_*L*_ ℙ (*F*)_*R*_.

The result of Step (*) is a vector of input states **I**_*x*_ and a vector of their probabilities **p**_**I***x*_.

**Step (**): computing outputs**. Step (**), the computation of the output states and probabilities given the input states and probabilities, is described by the following pseudocode:

I. generate possible output states from input states;
II. generate *Q*_*x*_, the Q matrix for node *x*, considering all possible input states and output states;
III. rearrange the order of input states to match the order of output states and construct a rearranged probability vector **p**_*x*_ from **p**_**I***x*_;
IV. compute output state probabilities using **p**_*x*_, *Q*_*x*_, and Eqn. 36;

To use Eqn. 36 to obtain output probabilities, the state space of **p**_*x*_ must include all possible output states. Thus, it is necessary to find all possible output states **O**_*x*_ for a set of input states **I**_*x*_.

(Step I) Possible output states consist of all states that are accessible from any number of transitions starting from the set of input states, and they include the input states themselves. The set of possible output states **O**_*x*_ is computed via a recursive algorithm that finds all states that are accessible through a one-step transition from the current set of states, and runs until all such transitions are already included in the set.

(Step II) Once the state space **O**_*x*_ is enumerated, the Q matrix can be constructed using the procedure described in Section A.1.

(Step III) As a minor technical point, to apply matrix operations, the input state vector **I**_*x*_ and the corresponding probabilities **p**_**I***x*_ must be rearranged to match the order of states enumerated in Step I, and an input probability of 0 must be assigned to the states in **O**_*x*_ that are not part of **I**_*x*_. The rearranged input probability vector is **p**_*x*_.

(Step IV) Once **p**_*x*_ is obtained, Eqn. 36 is used to compute the output state probabilities. The matrix exponential in our algorithm is computed by the function `linalg.expm` in the package `scipy`.

## Acknowledgments

We are pleased to contribute to the Mike Waterman special issue this application of a recursive algorithmic approach to a problem in coalescent theory and phylogenetics. We acknowledge support from NIH grant R01 GM131404.

## Notes

### Competing Interest Statement

The authors have declared no competing interest.

